# Divergent selection on behavioral and chemical traits contributes to isolation between populations of *Drosophila melanogaster*

**DOI:** 10.1101/2020.09.28.316703

**Authors:** Bozhou Jin, Daniel A. Barbash, Dean M Castillo

**Affiliations:** Department of Molecular Biology and Genetics, Cornell University, Ithaca NY; School of Biological Sciences, University of Utah, Salt Lake City, UT

**Keywords:** sexual selection, cuticular hydrocarbons, plasticity, speciation

## Abstract

Speciation is driven by traits that can act to prohibit mating between nascent lineages, including male courtship and female preference for male traits. Mating barriers involving these traits evolve quickly because there is strong selection on males and females to maximize reproductive success, and the tight co-evolution of mating interactions can lead to rapid diversification of sexual behavior. Using lineages of *D. melanogaster* that show strong asymmetrical reproductive isolation, we ask two key questions: which specific male traits are females selecting, and are these traits under divergent sexual selection? These questions have proven extremely challenging to answer, because even in closely related lineages males often differ in multiple traits related to mating behavior. We address these questions by estimating selection gradients for male courtship and cuticular hydrocarbons for two different female genotypes. We identify specific behaviors and particular cuticular hydrocarbons that are under divergent sexual selection and likely contribute to reproductive isolation. Additionally, we discovered that a subset of these traits are plastic; males adjust these traits based on the identity of the female genotype they interact with. These results suggest that even when male courtship is not fixed between lineages, ongoing selection can contribute to reproductive isolation.

## Introduction

Sexual selection is a powerful force that drives courtship behavior evolution in populations, and can eventually lead to reproductive isolation (Kirkpatrick 1982; Andersson 1994; Boughman 2002; Coyne and Orr 2004; Ritchie 2007). This observation is supported by comparative studies that quantify rapid trait evolution across widely diverged clades (Barraclough *et al.* 1995; Moller and Cuervo 1998) as well as by studies that estimate selection gradients on sexual traits in focal populations within species (Hill 1991; Brooks and Endler 2001; Rebarm *et al.* 2009; Callander *et al.* 2012; Oh and Shaw 2013; Steiger and Stokl 2014). Sexual selection can vary between populations, which could be a prerequisite for divergent selection and speciation (Chenoweth *et al.* 2010; Watts *et al.* 2019), but documenting ongoing divergent selection in the context of lineage divergence is more difficult (Higgie *et al.* 2000; Maan *et al.* 2004; Svensson *et al.* 2006; Maan *et al.* 2010; Pauers and Mckinnon 2012; Langerhans and Makowicz 2013; Selz *et al.* 2016; Wilkins *et al.* 2016). The difficulty in making connections between ongoing sexual selection and reproductive isolation may reflect that the most intensely studied traits for premating reproductive isolation are conspicuous fixed differences between well-established species (Mckinnon and Rundle 2002; Coyne and Orr 2004; Qvarnstrom *et al.* 2010). This contrasts with recently diverged species that may not show fixed differences in key mating-related traits, but nevertheless show strong reproductive isolation (Mallet 2008; Hendry *et al.* 2009; Merot *et al.* 2017; Khallaf *et al.* 2020a). By estimating selection gradients in lineages that experience gene flow yet show strong isolation, one can determine which traits contribute to reproductive isolation and whether ongoing selection reflects divergence in traits that contribute to reproductive isolation.

One reason why it is important to study ongoing (i.e. contemporary) sexual selection is because the targets or intensity of sexual selection may have changed over the course of speciation (Schluter and Price 1993; Price 1998), especially if mating signals are context- or environment-dependent (reviewed in Candolin 2019). When species differences are fixed, ongoing sexual selection can continue to act, but on mating traits not directly associated with reproductive isolation (Ryan and Rand 1993; Boake *et al.* 1997). In nascent lineages, however, one is much more likely to identify divergent selection on traits that contribute directly to reproductive isolation. One challenge for studying nascent lineages is that suites of traits show correlated divergence (Hohenlohe and Arnold 2010; Oh and Shaw 2013). When multiple correlated traits are divergent it can be difficult to disentangle which traits are important for female mate choice (Hohenlohe and Arnold 2010). This can be especially true when there are overlapping trait values between lineages, or if males have the potential to change their courtship based on female identity (Pfennig *et al.* 2010; Berdan *et al.* 2019; Fox *et al.* 2019).

In models of speciation by sexual selection, it is often assumed that there are single optimal male traits that are selected by a uniform female preference (Lande 1981; Kirkpatrick 1982; Kirkpatrick and Ravigne 2002). However, variation in female preference often occurs, where one male phenotype is not uniformly preferred (Jennions and Petrie 1997; Rebar and Rodriguez 2013; Mendelson *et al.* 2014; Kelley 2018). The maintenance of variation in male courtship traits and female preferences can lead to frequency-dependent selection (Otto *et al.* 2008), which in turn can lead to rapid evolution of reproductive isolation if populations become geographically isolated (Otto *et al.* 2008; Mendelson *et al.* 2014; Castillo and Delph 2016). Variation in female mate preference could also maintain selection for male courtship plasticity, if males maximize fitness by tailoring courtship to match female preference. Males can alter mating-related traits such as the intensity of courtship or size of ejaculate, based on the presence of rivals or based on the mating status of females (reviewed in Bretman *et al.* 2011; Petfield *et al.* 2005; Kelley and Jennions 2011; Otte *et al.* 2018). However, the potential for such behavioral plasticity to contribute to speciation has not been investigated.

In this study we use incompletely isolated lineages of *D. melanogaster* that show strong asymmetrical reproductive isolation to determine which traits are under divergent selection. *D. melanogaster* originated in southern Africa and migrated out of Africa in the past 10,000-15,000 years (Li and Stephan 2006; Pool *et al.* 2012; Kapopoulou *et al.* 2018). This resulted in lineages that remained in the ancestral range, called Z-type, and cosmopolitan lineages, M-type, that spread worldwide (Wu *et al.* 1995; Hollocher *et al.* 1997). Reproductive isolation between these lineages is asymmetric; M-type females mate indiscriminately when given a choice between males of each type, while Z-type females typically show strong preference for Z-type males (Wu *et al.* 1995; Hollocher *et al.* 1997). Given that female preference has evolved rapidly, it is assumed that male courtship has also rapidly evolved; however, divergence in male courtship has not been thoroughly tested, aside from the role of a few cuticular hydrocarbons (Grillet et al. 2012).

We estimated selection gradients on courtship behaviors and cuticular hydrocarbons for both M-type and Z-type females to determine if divergent selection is acting on specific traits. From previous work using common M-type lab strains, we had *a priori* predictions that the time spent producing vibratory courtship song, and the production of cuticular hydrocarbons 7-tricosene (7-C23) and cis-vaccenyl acetate (cVA), would be under positive selection in M-type lineages (Wu *et al.* 1995; Talyn and Dowse 2004; Grillet *et al.* 2006; Kurtovic *et al.* 2007; Scott *et al.* 2011). We predicted that these traits might be used for Z-type females to discriminate against M-type males (Grillet *et al.* 2012), similar to a species recognition trait. But to make connections between divergent selection and reproductive isolation, we needed to determine which traits are under selection specifically in the Z-type lineage.

## Methods

### Fly strains

All stocks were reared on standard yeast-glucose media prepared at Cornell University and kept at room temperature (~22°C). Information for each strain is in the Supplemental Information. For all mating experiments and cuticular hydrocarbon extractions, virgin males and females were collected and aged for 7-10 days. Each male fly was kept in an individual vial since male mating behavior is affected by their interactions with other males (Dixon *et al.* 2003).

### Video recording and analysis of courtship behavior

To record multiple courtship interactions simultaneously, we used a chamber design that allowed for easy loading of individuals with a mechanism to keep individuals separate until recording started (Koemans *et al.* 2017). All mating experiments were carried out within two hours of lights on (between 9:00AM and 11:00AM local time) as this is the time we could observe the maximum number of copulations. In pilot recordings we found no differences in courtship intensity or copulation rate for recordings that were done early or late in our recording window. The recording chamber was placed into a lightbox with LED strips (Konseen 16” Square Mini Dimmable Photo Light Box) to control light intensity across trials and courtship was recorded for 30 minutes with a camera (RoHS 0.3MP B&W Firefly MV USB 2.0 Camera). We used two courtship chambers to allow each chamber to be empty between recordings so any volatile cues from the flies would dissipate.

We used the software BORIS (Friard and Gamba 2016) to quantify male behavior. Behaviors were classified manually because we were interested in potentially identifying novel behaviors. We recorded all behaviors as state behaviors at 10 second intervals (similar to Gaertner *et al.* 2015). This might underestimate the time spent on a behavior, but still captures large differences in behavioral sequence since on average, interactions were 10.5 minutes long.

During the first stage of observations, we attempted to identify all behaviors that were occurring. Most were well described elements including singing, chasing, licking, attempted-copulation, copulation, and scissoring (Cobb *et al.* 1985; Cobb and Jallon 1990). We noticed that singing behavior occurred at two positions relative to the female, so we designated a separate singing (singing-2 is mostly from in front of the female, head-to-head, compared to “normal” singing with the male behind or slightly to the side of the female). We also noticed several “circling” behaviors where the male moved in an arc around the female while wing displaying (singing or scissoring; Supplemental Video 1). The frequency of circling behaviors across this initial dataset indicated that only one circling behavior was frequent enough to reliably identify, so all circling behaviors were combined into a single metric. After watching many videos we could not reliably score licking and tapping, so we combined these with chasing and following behaviors into “engaging”. This left 8 categories: separate, engaging, singing, singing-2, scissoring, circling, attempted-copulation, and copulating.

When scoring the male behaviors, the video was run at 2X speed and paused every 10 seconds to record a behavior. The observer recorded observations blindly with respect to genotype. The scoring for trials that successfully copulated were ended as soon as the flies started copulation. There were several trials that were unsuccessful in mating that we watched to make specific comparisons (see below in Statistical analysis). These unsuccessful trials were quantified for a time equal to the average copulation latency of the successful trials for the same genotype combinations.

### Cataloging potential Z-type behaviors

Though the courtship sequence of *D. melanogaster* is well known, this information comes exclusively from M-type strains (Cobb *et al.* 1985) so it was critical to establish what, if any, additional behaviors may occur in Z-type strains. The *D. simulans* strain served as a standard for the scissoring behavior as it is a frequent part of their courtship display (Cobb *et al.* 1985; Supplemental Video 2) and some genotypes of *D. melanogaster* with African ancestry may engage in this behavior (Yukilevich and True 2008). We recorded six pairs of flies during every trial block. This balanced efficiency with the need to record pairs at a close distance to document subtle behaviors.

### Selection acting on courtship behaviors

Our second stage of recording was aimed at identifying behaviors under selection by correlating mating behaviors with copulation success and copulation latency, which was our proxy for mating success and fitness (Hoffmann 1999; Taylor *et al.* 2007). We conducted trials between Z53 or DGRP882 females and the males from our panel of M and Z-type genotypes. During each block, ten pairs of flies were video recorded simultaneously with five pairs courting Z53 females and the remaining five courting DGRP882 females. We used five male genotypes per block such that the Z53 and DGRP882 females were paired with the same genotypes within a block, with the male genotypes determined from a randomized sequence of five. The order of the sequence was rotated after every three blocks of recording, so that each strain had equal probability of being in an early vs late block across the course of the experiments. This randomization scheme limited the effects on courtship intensity of time of day and of blocks occurring across multiple weeks. We repeated this scheme until all genotype combinations had five successful matings, which resulted in some combinations having more successful matings than others.

### Cuticular hydrocarbon collection and quantification

We collected cuticular hydrocarbons (CHCs) from three males and three females for all of the genotypes used in the behavioral studies, except for the *D. simulans* strain. CHCs are influenced by both temperature and humidity so ensured that all collections were done in parallel to the behavioral observations (Noorman and Den Otter 2002; Savarit and Ferveur 2002). Male and female virgins were collected, stored individually and aged 7 days before CHCs were extracted in 8 dram glass vials with 100μl HPLC grade hexane (99% purity) for 1 min on an orbital shaker. The solvent was then transferred into an autosampler vial with a small volume insert, left in a fume hood for four hours to evaporate, and vials sealed and placed at −20°C until analysis. For analysis CHCs were re-eluted in 50μl hexane with an internal standard (hexacosane C27) based on (Dembeck *et al.* 2015).

To measure plasticity in male CHCs as a product of interacting with different female genotypes, we exposed Z53 and DGRP882 males to females from their own and opposite genotypes, while keeping additional virgin males with no contact as controls. After allowing courtship for 10 min we separated the pairs and placed males in individual vials for 30 min followed by immediate CHC extraction, hoping that males would be exposed to females but would avoid physical contact via copulation. Within the 10 minutes, however, all of the flies had copulated except DGRP882 males paired with Z53 males. We did not find however, known “female” CHCs on the males, confirming that any changes to the males were not the result of female CHCs rubbing off of males (Khallaf *et al.* 2020b). Quantification of CHCs is described in the Supplemental Information.

### Statistical analyses

#### Summarizing courtship behavior

To determine copulation latency and the percentage of time spent on each behavior we needed to estimate courtship initiation. We estimated courtship initiation in two ways: the time when the first non “separate” behavior took place or the time after three consecutive non “separate” behaviors took place. The rationale behind the second method was to account for observations where males and females were close and scored as “engaging” but courtship was broken off and there was a long gap before courtship was reinitiated. For the majority of the cases the difference between courtship initiation based on the two methods was minimal (mean=7.4 seconds) and did not influence the overall estimate of copulation latency. For a few observations there was up to 4 minutes difference (max=262.318 seconds) and re-evaluating these outliers it was clear that the first “engaging” was not courtship initiation. Given this comparison, we used the second method as it was more accurate. The percentage of time spent in a particular behavior was calculated as the number of timepoints scored as this behavior, divided by the total number of timepoints recorded until copulation occurred. We determined that courtship was similar across recording periods and determined when copulation failure was a result of males not initiating courtship (Supplemental Information).

#### Mating rejection in long-term experiments

After analyzing videos we observed that there were several genotype combinations that failed to copulate. We expected Z53 to reject known M-type males; however there were several genotype combinations that we did not a priori expect to fail, including Z53 females x LZ21 males and DGRP882 females with Z30 or LZ21 males. To determine if this was a function of our short recording time we set up single-pair matings between these genotypes and measured the time until progeny were first seen over a 16 day period (Supplemental Information).

#### Plasticity analysis

To determine if male trait values for both behavioral traits and CHCs changed with respect to which female genotype a male was courting, we looked for evidence of consistent phenotypic plasticity across genotypes. We evaluated models that included male and female genotype and their interaction on each trait. The male genotype effect would capture differences between males. The female genotype effect would indicate plasticity if males, on average, exhibited different trait values when interacting with different females. This pattern would graphically appear as parallel reaction norms (Roff 1997). We included the interaction effect into our models because this would demonstrate that males differed in their effects and did not have consistent plasticity. To test these effects we used aligned ranks transformation (ART) ANOVAs using the ARtools package in R (Kay and Wobbrock 2019). Details of these models and rationale for using them are included in the Supplemental Information.

We also tested for plasticity in how the courtship behavioral sequence was carried out by males when interacting with different females. We quantified this difference using transitions between behavioral states. Behavioral sequences can be viewed as Markov chains, and analyzing transition matrices in this Markov chain framework can determine differences in a behavior sequence in courtship behavior (Markow 1987; Gaertner *et al.* 2015). First we constructed a transition matrix by summing all of the transitions across all trials for a specific female genotype x male genotype combination. Then we tested whether the transition matrix for a given male genotype differed when interacting with the different female genotypes, using a homogeneity test from the Markov package in R (Spedicato 2017). If these matrices were not homogeneous we could infer that the overall transition rates were different but not which particular transitions were responsible for this pattern. To determine if there were consistent transition differences we looked for plasticity of individual transitions after a variable reduction process using the VSURF package in R (Genuer *et al.* 2019).

For the CHC plasticity analysis we used dimension reduction to avoid multiple testing issues. While principal components (PC) are a convenient way to summarize this type of multivariate data, PC scores suffer from the inability to examine/interpret the effects of individual variables. Instead we used the same random forest procedure that we used for the transition data. For this classification problem CHCs could be explained by a categorical variable that captured the interaction of male x female genotype (6 levels for this experiment, see methods above). From the remaining variables we carried out ART ANOVA testing for male, female and male x female genotype effects.

#### Detecting selection acting on male behavioral traits

To determine if directional selection was acting on particular male traits and whether it was acting in the same direction across the two female genotypes, we used quantile regression to determine relationships between the male trait and copulation latency (our proxy for mating success and fitness). We used quantile regression to condition our regression model on the median rather than the mean, which is useful for non-normally distributed data (Koenker 2005). We used the package quantreg in R (Koenker 2019) to fit models that included effects of female genotype, the trait of interest, and the interaction of genotype and trait on copulation latency. The effect of female genotype often captured the fact that the distribution of traits was not identical between female genotypes. One reason was plasticity (see results) the other was that not all genotypes successfully copulated with both females. As a result the same male genotypes were not present for both female genotypes. The effect of the continuous trait on copulation latency was a slope and can be interpreted as the selection gradient. The interaction term is the change in slope/selection gradient that occurs when interacting with the second female genotype. Because we did not measure CHCs for each male used in the courtship trials, we used the mean to represent the CHCs of male genotypes. To be consistent we correspondingly used the mean for male behavioral traits, which was also necessary given the differences in sample sizes of successful trials.

For behavioral traits we used quantile regression on engaging, circling, and attempted-copulation. Because singing-2 was infrequent and correlated with time spent singing, we combined these into a single metric. The courtship behaviors were not strongly correlated within a male genotype, or between female genotypes so we analyzed each trait independently (Supplemental Table 1). For the analysis of CHCs on copulation latency we used random forest to reduce the number of variables before completing regression. The VSURF package only allows for a single effect that is either continuous or categorical, which precluded variable selection on our full model of interest. We therefore examined predictive relationships between the copulation latency and the amount of a specific CHC compound using the complete dataset (copulation latency across all genotype combinations), and reduced datasets that represented the individual female genotypes. In this way we expected to identify CHCs that might be important to one, but not the other female genotype. We then estimated the selection gradients and reported confidence intervals for each parameter, to demonstrate how/if the selection gradient changes between female genotypes. We also included the compounds cis-vaccenyl acetate (cVA) and 7-tricosene (7-C23) because of their reported importance to mating interactions in *D. melanogaster* (Grillet *et al.* 2012), but these were not identified as important to selection from the random forest variable reduction.

For a subset of behavioral traits and CHCs we also wanted to determine if selection was acting on the probability of mating occurring, rather than on the speed by which it occurred. For this analysis we used binomial regression with the number of successful and unsuccessful copulations per female genotype x male genotype combination as the dependent variable. We focused on the scissoring and circling behaviors since they did not show a linear response for the Z-type female genotype (see results), but appear to be Z-type specific behaviors. We also included singing since this is thought to dominate M-type courtship (Cobb *et al.* 1985; Talyn and Dowse 2004). For CHCs we focused on 7-C23 and 5-tricosense (5-C32), because they are involved in mating probability (Grillet *et al.* 2012) as well as cVA.

#### Correlations Between Courtship Traits and Cuticular Hydrocarbons

Females that strongly prefer Z-type over M-type males often have a distinct CHC profile that is potentially dependent on their *desat2* alleles (Takahashi *et al.* 2001; Fang *et al.* 2002; Chertemps *et al.* 2006; Coyne and Elwyn 2006a). Z-type female preference and male acceptance by a Z-type female are expected to be correlated (Hollocher *et al.* 1997). To determine if there were correlated traits that are diagnostic of Z-type males we first estimated the correlation between phenotypic distance of CHCs for male and female genotypes, assuming that the presence of the Z-specific compound 5,9-heptacosadiene (5,9-HD) is diagnostic of Z-type female. We then used principal component analysis (PCA) on the combined male courtship and behavior data to identify strongly correlated traits.

## Results

### Courtship of Z-type and M-type males is significantly different

We quantified the behavior of a standard *D. melanogaster* M-type strain (DGRP882), a typical Z-type strain (Z53), and one strain of *D. simulans* (SA22) to identify similarities and differences among these lineages. To our knowledge, all previous studies of male courtship behavior in *D. melanogaster* used only M-type strains. We identified several behaviors in the Z-type strain that did not occur in the M-type strain, but do occur in other species of the *melanogaster* subgroup. This included scissoring, where males open and close their wings rapidly (Supplemental Video 2), similar to scissoring found in *D. simulans* (as defined in Cobb *et al.* 1985). We also observed circling, where males move around the female in an arc while vibrating or scissoring at the same time (Cobb *et al.* 1985; Supplemental Video 1). In addition, the overall time spent on behaviors common between the two *D. melanogaster* genotypes was significantly different. Specifically, the Z-type strain spent less time singing compared to the M-type strain, but more time singing than *D. simulans* (Supplemental Fig. 3). For circling, we did not observe this behavior in the M-type strains but there was ample variation across Z-type strains.

### Z-type and M-type strains are reproductively isolated

Despite active male courting, Z53 females strongly rejected both DGRP882 and Canton-S M-type males, consistent with previous reports of asymmetric reproductive isolation in this system (Wu *et al.* 1995). This rejection persisted much longer than the limited time we video recorded. To expand our ability to assay isolation between strains, we used progeny production as a proxy for mating success. Comparing control matings between Z53 females x Z53 males with matings between Z53 females and several other genotypes, we saw that larvae took significantly longer to appear in crosses between Z53 females and the M-type males DGRP882 and Canton S as well as for ZH33 males (Supplemental Fig. 4A). . Due to the small sample sizes and crossing survival curves, we could not determine if these male genotypes were significantly different from each other, but qualitatively fewer crosses produced larvae when M-type males (DGRP882 and Canton S) were paired with Z53 females (Supplemental Figure 4A).

Based on the literature we did not expect reproductive isolation between M-type females and Z-type males (Wu *et al.* 1995; Coyne and Elwyn 2006). Nevertheless, some genotypes did not mate within the 30 minute video recordings, so we again assayed for progeny production. For crosses with DGRP882 females we used ZH33 males as our baseline because they quickly produced both eggs and larvae indicating that they mated within the first 24 hours (Supplemental Fig. 4B). Compared to ZH33 males, LZ21 and Z30 males took significantly longer to produce larvae and in fact very few matings were successful over the course of the 16 day experiment (Supplemental Fig. 4B). This level of reproductive isolation was similar in magnitude to the Z53 female x M-type male crosses. We conclude that isolation is not fully asymmetric and that males contain significant variation in mating behavior.

### Male courtship behavior is plastic

Having found significant differences between the courtship of Z53 and DGRP882 males, we next tested whether male courtship is a static or plastic trait. Using a panel of Z-type and M-type males we quantified the differences in behaviors when males interacted with either a Z- or M-type female. For the majority of the male genotypes, the time spent carrying out specific behavioral traits was female-genotype dependent. The two traits that showed consistent plasticity were attempted-copulation and singing (Supplemental Table 2). The strongest signal was for singing, where males consistently spent more time singing when courting the M-type female (Figure 1). Engaging and circling also had significant genotype effects but there was not a consistent pattern of plasticity across genotypes (Supplemental Table 1).

**Figure 1.**
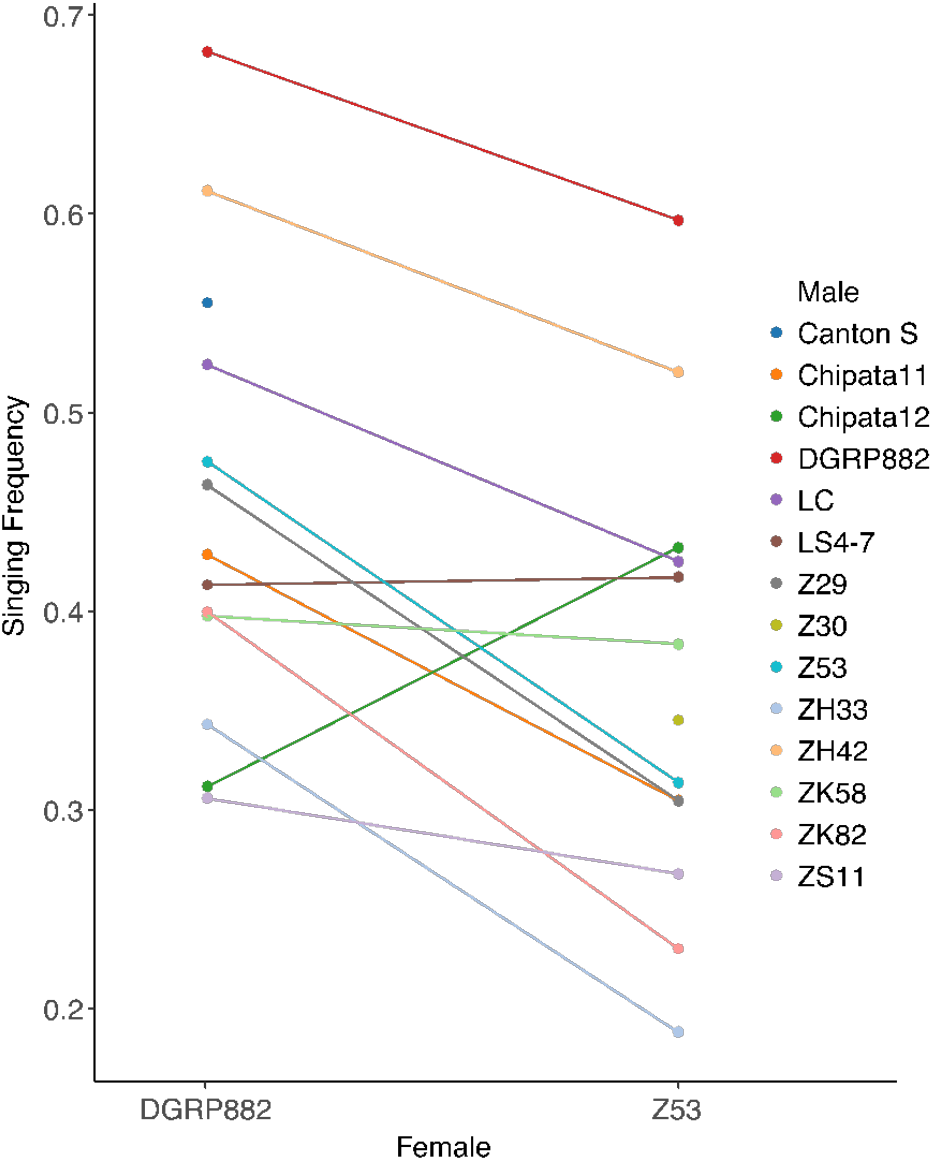
Male courtship singing frequency is female-genotype dependent. There was a consistent decrease in singing frequency for most strains when males of each genotype were presented to a Z53 female, compared with being presented to a DGRP-882 female. Each point represents the average singing frequency of the given male genotype (indicated by color) when presented to a female of either genotype. Points that occur only in Z53 or DGRP882 female backgrounds indicate that males did not copulate with the other genotype. This includes Canton S males only copulating with DGRP882 females and Z30 males only copulating with Z53 females. DGRP882 male behavior was analyzed when courting Z53 females even though they did not copulate.

### Courtship plasticity changes the behavioral sequence

Given that the time spent on behaviors was plastic, this could affect the behavioral sequence and transitions between behaviors. We therefore compared transition matrices for male genotypes interacting separately with the two female genotypes. Seven out of the twelve male genotypes had significantly different transition matrices (Supplemental Table 3). The transitions that contributed to this plasticity were singing-attempted-copulation and engaging-singing (Supplemental Table 4). When interacting with Z-type females the male genotypes transitioned more from singing to attempted-copulation and transitioned less from engaging to singing (Supplemental Fig. 5). This reflects that males were singing less in this interaction. The transition between singing-scissoring had significant differences between male genotypes, but was not female-genotype dependent (Supplemental Table 4).

### Male cuticular hydrocarbons are plastic

We next tested the hypothesis that males change their CHCs upon exposure to females by quantifying CHCs in Z53 and DGRP882 males that had either been kept virgin or were exposed to Z53 and DGRP882 females. After variable reduction we were left with five compounds from the original twenty-one that we had identified, four of which had significant genotype and treatment effects (Supplemental Table 5). The compounds 5-pentacosene (5-C25) and 9-pentacosene (9-C25) showed similar patterns, with large differences between DGRP882 and Z53 males in all treatments (Supplemental Fig. 6). Z53 males had reductions in 5-C25 when exposed to either female strain (Supplemental Fig. 6). The other two compounds had changes that depended on both the male and female genotype. Males reduced their amount of the compound 7-tricosene (7-C23) when they were exposed to females of the opposite type. For 2-methyltriacontane (2-Me-C30), DGRP882 males increased their amount of this compound with DGRP882 females, but Z53 males decreased this compound with DGRP882 females (Supplemental Fig. 6). The plastic changes may contribute to selection on CHCs in terms of mating success (see below).

### Divergent selection acts on courtship traits

To test for directional selection on courtship traits within a lineage, and divergent selection between lineages, we used quantile regression to determine the relationships between the six behavioral traits and copulation latency. We assumed shorter courtship latency was a proxy for increased mating success. A negative regression coefficient indicates a positive selection gradient and a trait that is favored by a specific female genotype. Attempted-copulation was the only trait that had divergent selection gradients for the two female genotypes. There was a significant relationship between increased attempted-copulation and shorter copulation latency (positive selection gradient) for Z53 females, and the opposite pattern for DGRP882 females (Figure 2A; Supplemental Table 6). For singing we only observed selection in one female background, with no relationship for DGRP882 females, but a significant relationship between more singing and longer copulation latency (negative selection gradient) for Z53 females (Figure 2C; Supplemental Table 6). Lastly, for both genotypes there was a similar relationship between more scissoring and longer copulation latency (Fig. 2B; Supplemental Table 6).

**Figure 2.**
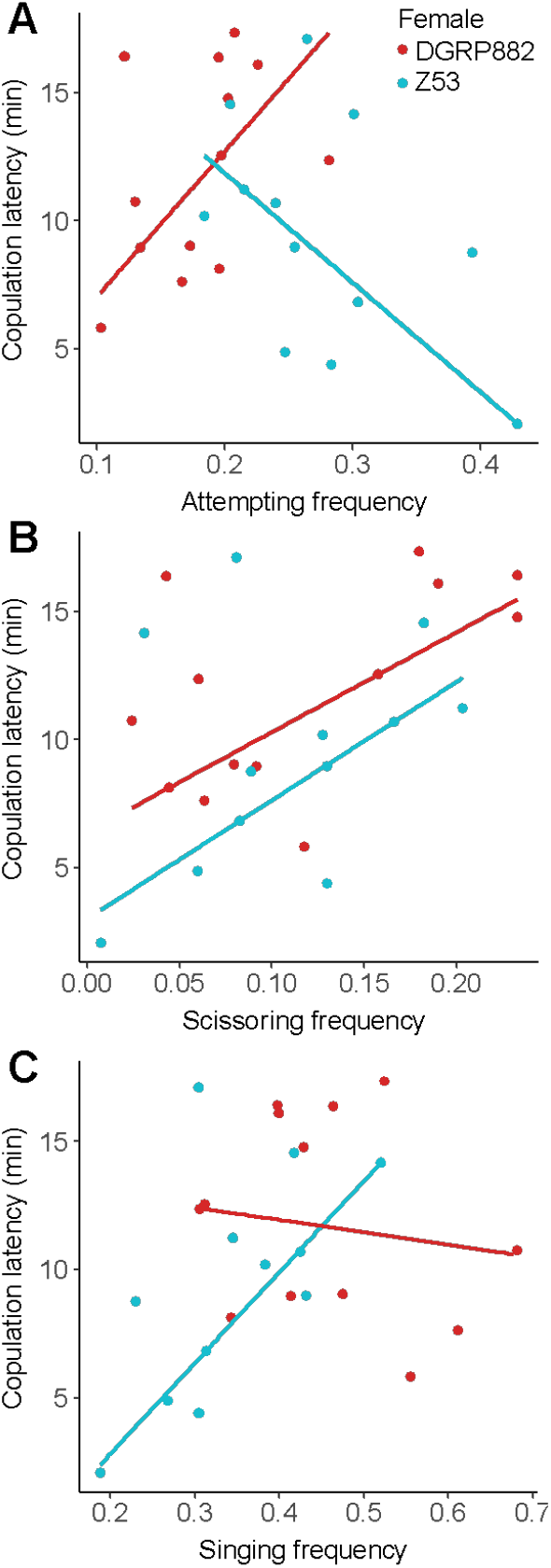
Directional selection on male courtship behavior is female-genotype-dependent. A) There was a negative relationship between attempted-copulation frequency and copulation latency for males interacting with Z53 females and the opposite pattern for males interacting with DGRP-882 females. B) There was a similar positive relationship between scissoring frequency and copulation latency for DGRP-882 and Z53 females. C) There was a positive relationship between singing frequency and copulation latency for Z53 females but no relationship for DGRP-882 females. Shorter copulation latency is a proxy for increased mating success. Each point in the figures represents the average behavioral frequency and copulation latency of a male genotype when presented to a female of the given genotype. The regression lines were produced using quantile regression.

In addition to testing for linear relationships we determined if courtship traits predicted whether a female would copulate (successes vs failures) using binomial regression. We analyzed the effects of singing, scissoring, and circling on the probability of copulation. This type of selection can occur if there is a threshold for acceptance of mating. For singing, the pattern for Z53 females was similar to the linear selection analysis, where more singing decreased the probability of copulation (Supplemental Table 7). In this analysis we did observe divergent selection for DGRP882 females, where more singing increased the probability of mating (Supplemental Table 7). The time spent scissoring was not related to the probability of copulation for DGRP882 females, but did increase the probability of copulation for Z53 females (Supplemental Table 7). The time spent circling did not have significant effects on the probability of copulation for either female genotype.

### Divergent selection acts on cuticular hydrocarbons

We next estimated directional selection of cuticular hydrocarbons (CHCs) to test which compounds contributed to mating success, and if there were differences between the female genotypes. After variable reduction we reduced the number of CHCs we examined from twenty-one to five compounds. We included two additional compounds cis-vaccenyl acetate (cVA) and 7-tricosene (7-C23) based on *a priori* expectations of their impact on mating success (see Methods). Four compounds had divergent selection gradients that were female-genotype dependent. For tetracosane (n-C24), 9-pentacosene (9-C25), and 2-methyl-hexacosane (2-Me-C26) there was a significant relationship between greater quantities of these compounds and decreased copulation latency for Z53 females and an increased copulation latency for DGRP882 females (Figure 3; Supplemental Table 8). For cVA the opposite pattern was true, greater quantities reduced copulation latency for DGRP882 females and increased copulation latency for Z53 females (Figure 3D; Supplemental Table 8). The remaining two compounds did not have significant relationships with copulation latency.

**Figure 3.**
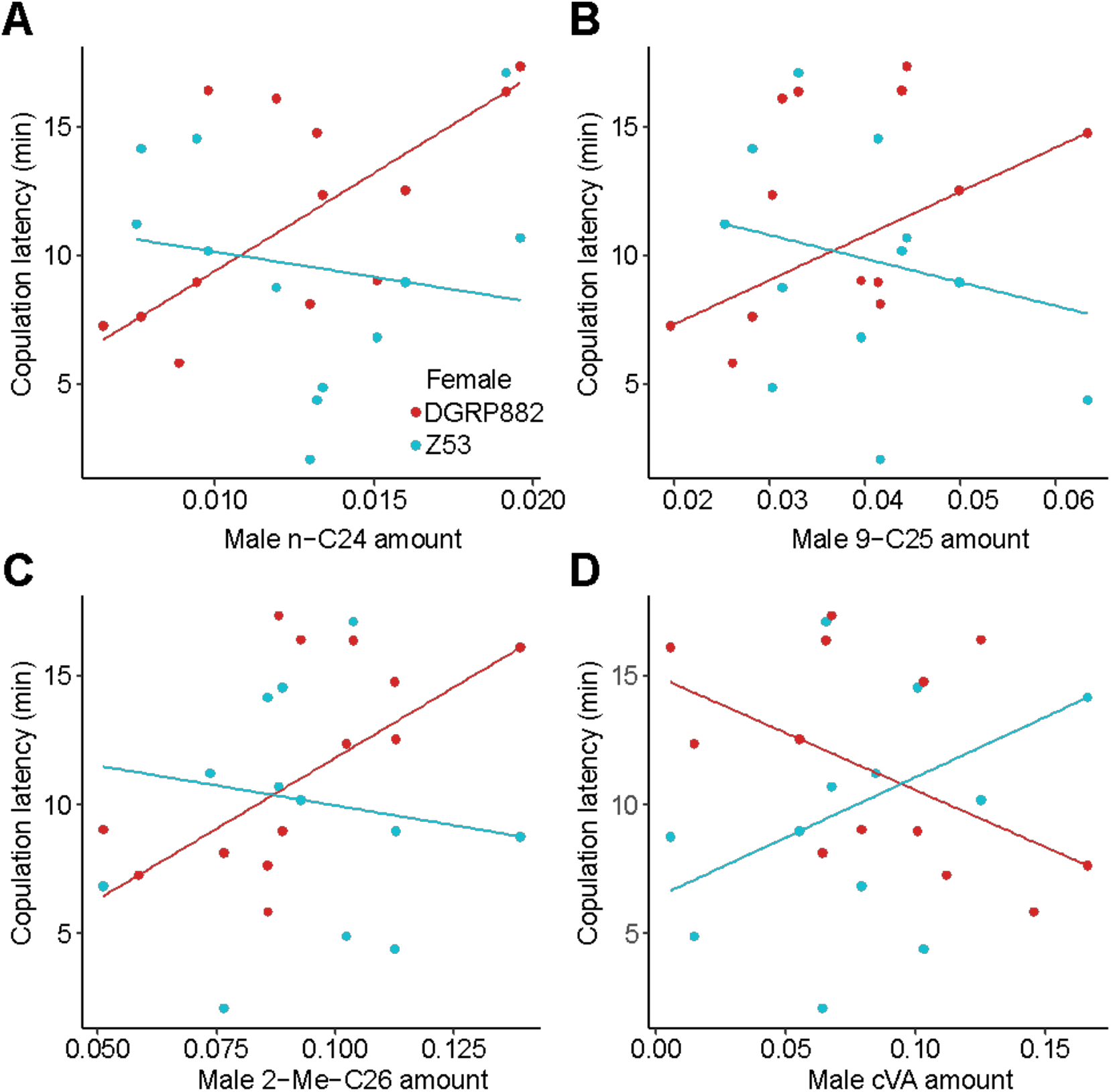
Directional selection on male cuticular hydrocarbons (CHCs) is female-genotype-dependent. There were significant positive relationships for DGRP-882 females and negative relationships for Z53 females for copulation latency and the amounts of three CHCs in males including A) tetracosane, n-C24, B) 9-pentacosene, 9-C25 and, C) 2-methyl-hexacosane, 2-Me-C26. D) The opposite pattern occurred for cVA with a negative relationship between the amount of cVA and copulation latency for males with DGRP-882 females and a positive relationship with Z53 females. Shorter copulation latency is a proxy of increased mating success. Each point in the figures represents the average relative abundance of the corresponding CHC and copulation latency of a specific male genotype, when presented to the indicated female genotype. The regression lines were produced using quantile regression.

We used binomial regression to test the hypothesis that the amounts of 7-C23, 5-C23, and/or tricosane (n-C23) predict probability of copulation. These compounds had no effect on copulation probability for DGRP882 females, but 7-C23 and n-C23 had significant effects on decreasing copulation probability for Z53 females (Supplemental Table 9). When we analyzed cVA in binomial regression the results were consistent with the linear regression. Greater quantities of cVA increased mating probability with DGRP882 females and reduced mating probability with Z53 females (Supplemental Table 9).

### Variation in courtship traits results in discrete clusters of Z-type male genotypes

Given the behavioral trait differences between the male genotypes and their impacts on mating success, we wanted to determine if there was a suite of traits and CHCs that together could be diagnostic of a Z-type male. Most of our prior information on Z-type flies comes from female choice and CHC experiments, so we first determined the distributions of the C27 isomers 7,11-heptacosadiene (7,11-HD) and 5,9-heptacosadiene (5,9-HD) in our strains. The distributions of these compounds had large variation, separating the two known M-type strains (DGRP882 and Canton S) and ZH42 from the other Z-type strains (Supplemental Fig. 7). The M-type strains had high quantities of 7,11-HD and low quantities of 5,9-HD. Most of the suspected Z-type genotypes produced less 7,11-HD than the M-type strains, but there was a large range in the quantity of 5,9-HD produced by these genotypes (Supplemental Fig. 7). We determined if there was a phenotypic correlation between CHCs of males and females using Euclidean distances with respect to the Z53 strain (*r*=0.561, *P=*0.044). This captures the overall CHC differences in genotypes without relying on a single compound. Z53 males and females both had the largest phenotypic distance from DGRP882, and across all genotypes the CHC phenotypic distances were greater for female compounds than male compounds. The correlation was not as strong as might have been expected though, because of genotypes that are divergent from both DGRP882 and Z53 in phenotypic space (Supplemental Fig. 8).

Given this variation in CHC phenotypes we determined using principal component (PC) analysis the relationships between male CHCs and male behavioral traits when courting Z53 females. While genotypes were spread across the PC space, there was clustering based on the loadings of specific combinations of behaviors and CHCs (Figure 4). For example, one cluster included ZH42 and DGRP882 and the loadings that contributed were Singing, cVA and 7-C23. In the opposite direction in PC space, Z53 and a few other Z-type strains clustered with contributing loadings including Attempted-Copulation and C25 isomers. The last main cluster included Z30, LS4-7, and other strains that were mostly separated by the Scissoring behavior and additional CHC compounds. Overall, the different clusters suggest there is not a single courtship and CHC combination that can diagnose all Z-type males.

**Figure 4.**
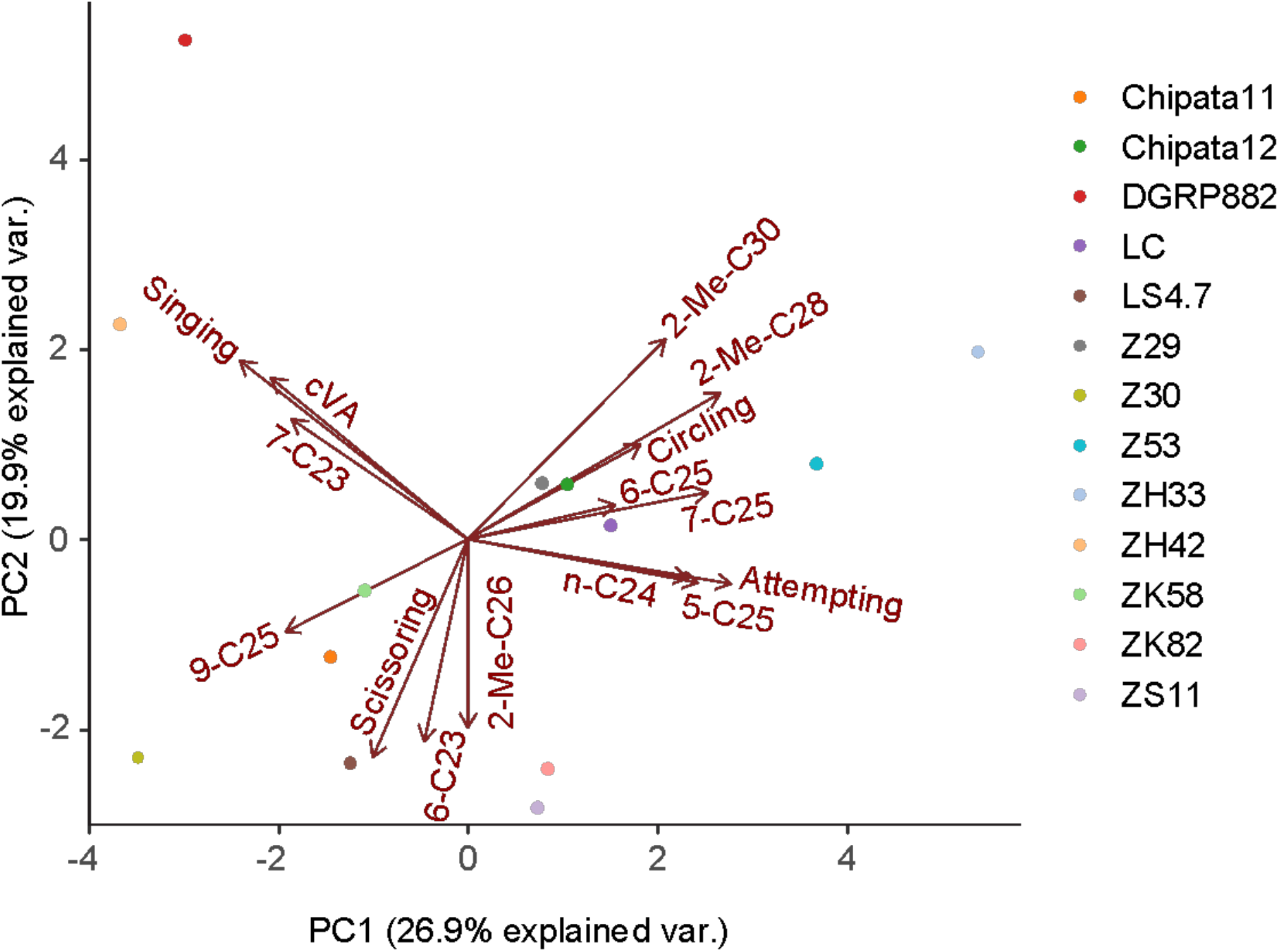
The combination of male cuticular hydrocarbons and behaviors that characterizes distinct male courtship phenotypes. The combination of singing behavior and the cuticular hydrocarbons cVA and 7-tricosene (7-C23) defines M-type genotypes. The presumably Z-type males fall into two broad clusters defined by unique behavior and CHC combinations. Each point represents the average value for a given male genotype with behavioral data coming from interactions with Z53 females. Each arrow represents one main variable used to construct the principal components, with its length representing the loading on a particular principal component axis.

## Discussion

While sexual selection appears to be an important driver of speciation based on macroevolutionary patterns, connecting contemporary divergent sexual selection to reproductive isolation has been challenging. We leveraged *D. melanogaster* to quantify selection on relevant male courtship traits for females from two diverging lineages. We comprehensively documented differences in male behavior and cuticular hydrocarbons (CHCs) between the M-type and Z-type lineages that allowed us to test for selection on these suites of traits. We confirmed that the time spent singing and the amount of the hydrocarbon cVA are under positive selection in M-type lineages. We then showed that these traits are under negative selection in Z-type lineages, where increasing time spent singing or increased amounts of cVA increased copulation latency. These findings identify these traits, as potential contributors to reproductive isolation. Interestingly, even though M-type strains are thought to mate indiscriminately, we identified traits under negative selection in this lineage. Together our results demonstrate that in early divergent lineages, there are direct connections between sexual selection and reproductive isolation, and that variation in courtship traits can be maintained by both male plasticity and variation in female preference.

Female mate choice is often correlated with the presence of a conspicuous fixed difference between species (Mckinnon and Rundle 2002; Coyne and Orr 2004; Qvarnstrom *et al.* 2010), but in young divergent lineages, trait variation may segregate between populations (Mallet 2008; Hendry *et al.* 2009; Merot *et al.* 2017; Khallaf *et al.* 2020a). As a result, both quantitative differences and the presence or absence of traits could shape reproductive isolation. There was large phenotypic variation among the African strains in our experiment, and for both courtship behavior and CHCs, we identified examples of divergent selection between lineages or selection within a single lineage. Given the recent and historical gene flow of European ancestry back into Africa (Pool *et al.* 2012), one might expect some of the variation to be due to this gene flow, with males phenotypically appearing either similar to M-type males or intermediate in phenotype. Indeed, we saw this with ZH42, which was similar in both behavior and CHCs to the M-type strains, but most Z-type strains were very distinct from M-type strains, suggesting ample variation segregating within African populations (Fig. 4). Interestingly, the traits under selection that we identified generally occur in both lineages.

We discovered that Z-type strains carry out two behaviors, scissoring and circling, that we did not see in M-type strains. Circling was not under selection in either lineage, but scissoring had a more interesting pattern. Scissoring has previously been observed in *D. melanogaster* strains from the Bahamas and Caribbean that contain African ancestry, but it was a minor part of their courtship (Yukilevich and True 2008). We found Z-type strains that display scissoring behavior at a higher frequency. Increased time spent scissoring increased the probability of copulation with Z-type females, but was under negative selection for both M-type and Z-type females. In fact, we saw very strong reproductive isolation between females from the M-type strain and males from the Z-type strain that displayed the most scissoring. This level of reproductive isolation was as strong as the isolation that is typical between Z-type females and M-type males (Supplemental Fig. 4). Scissoring thus provides an interesting example where a lineage-specific trait may contribute to reproductive isolation and shape female mate preference, but is not under divergent linear selection.

Both sexual selection and environmental selection can shape patterns of CHC divergence (Higgie *et al.* 2000; Greenberg *et al.* 2003; Chung *et al.* 2014), and are examples of magic traits, contributing to both ecological divergence and reproductive isolation (Servedio *et al.* 2011). CHCs strongly contribute to reproductive isolation in newly diverged lineages and often these differences are quantitative (Veltsos *et al.* 2012; Chung *et al.* 2014; Seeholzer *et al.* 2018; Khallaf *et al.* 2020a). By demonstrating divergent selection acting on CHCs we are able to determine how these quantitative differences contribute to reproductive isolation. We identified several CHCs that were under divergent sexual selection between lineages by focusing on African and Z-type strains and using data reduction techniques that do not rely on a priori candidate compounds. Previous studies did not determine which CHCs are under positive selection in Z-type lineages, which could be due to the phenotypic distribution of the genotypes in those studies and the focus on compounds that are typically in high quantities in M-type genotypes, specifically isomers of tricosene (Grillet *et al.* 2012). For the intensively studied compound 7-tricosene (7-C23) we recapitulated previous reports that it is under negative selection by Z-type strains (Grillet *et al.* 2012), but could not detect positive selection for M-type strains. One possibility is that all strains had the minimum amount of 7-C23 that would ensure acceptance by M-type females. If there is a threshold for mating acceptance based on this compound, it would be consistent with the idea that M-type females mate indiscriminately (Wu *et al.* 1995).

Another mechanism that could contribute to the asymmetry of reproductive isolation in this system is that male courtship is female-genotype dependent. Male courtship is described in seemingly contradictory terms, sometimes as a stereotypical sequence (Cobb *et al.* 1985; Gaertner *et al.* 2015), but other times as plastic (Dukas and Dukas 2012; Arbuthnott *et al.* 2017; Filice *et al.* 2020). This discrepancy might come from the fact that plasticity is often quantified in terms of courtship effort and intensity, rather than the behavioral sequence. We identified behaviors that were consistently plastic across genotypes and resulted in changes to the fundamental behavioral sequence. We also detected significant plasticity for some CHCs in *D. melanogaster,* which can be further explored in future studies. Plasticity that is female-genotype dependent has the potential to impact how we view sexual selection, by creating scenarios where males can maintain fitness in the presence of variation in female mate preference. Selection can maintain plasticity in recently diverged lineages where males come into contact with multiple female genotypes and gain fitness by adjusting their courtship displays. Asymmetrical reproductive isolation, which is especially common in *Drosophila* (Yukilevich 2012) and other systems where both females and males contribute to mate choice, might also reflect plasticity. For example, even though M-type males, such as DGRP882, reduced the time spent singing when they courted Z-type females, they still spent significantly more time singing than the Z-type genotypes and ultimately failed to mate with Z-type females. Female-genotype-dependent courtship plasticity suggests that males require lineage-specific female cues for successful courtship (Barker 1967; Cobb and Jallon 1990; Seeholzer *et al.* 2018). CHCs can both stimulate and inhibit mating (Ferveur 2005; Grillet *et al.* 2006; Ejima *et al.* 2007; Khallaf *et al.* 2020a; Khallaf *et al.* 2020b), and a strong difference exists in the female CHCs of Z-type and M-type females (COYNE *et al.* 1999; Dallerac *et al.* 2000). The ability of males to change their courtship can alter their courtship in subsequent mating interactions, which are typically not considered in the study of speciation.

The interaction between plasticity and learning can have important impacts on the strength of reproductive isolation and outcomes for speciation (Kujtan and Dukas 2009; Verzijden *et al.* 2012). Successful mating can create positive associative learning (Ejima *et al.* 2007; Griffith and Ejima 2009; Koemans *et al.* 2017; Zer-Krispil *et al.* 2018). If males adopt a successful courting strategy as a result of this conditioned learning they may show less plasticity in subsequent mating bouts (Dukas and Dukas 2012). If instead, plasticity is maintained across subsequent mating encounters, then gene flow would continue and reproductive isolation would not evolve (Kirkpatrick and Nuismer 2004; Pfennig *et al.* 2010). Plasticity within a species could therefore maintain variation in male courtship and female preference would persist in populations. This segregating variation could potentially facilitate rapid speciation, but only after geographic separation (Mendelson *et al.* 2014; Castillo and Delph 2016). We identified large differences in male behaviors and CHCs among Z-type strains, such that we could not define a single behavioral trait and CHC combination that classifies Z-type vs M-type males. This intriguing result suggests that there are segregating preferences in these populations. This variation, similar to what we currently observe, may have facilitated the rapid divergence of M-type strains from Z-type ancestors when these lineages migrated out of Africa. This has now created strong asymmetric reproductive isolation that is maintained by ongoing divergent sexual selection.

