## Supplemental Information for "Divergent selection on behavioral and chemical traits contributes to isolation between populations of *Drosophila melanogaster*"

***Fly Strains***

The *D. simulans* strain SA22 was provided by Chip Aquadro (originally collected in Stellenbosch South Africa). The *D. melanogaster* M-type strains were Canton-S (line maintained by Mariana Wolfner) and DGRP-882 (BDSC #28255). This DGRP line was chosen based on less African ancestry compared to other DGRP strains (POOL 2015). The remaining lines were chosen based on either prior knowledge of Z-type female behavior or collection location, with the goal of capturing as much variation in Z-type male behavior as possible. Whether a strain is Z-type or M-type has historically been classified by female preference behavior using choice tests, where females are presented with a choice between a M-type and Z-type male. Z-type females choose Z-type males preferentially (WU *et al.* 1995; HOLLOCHER *et al.* 1997). There is some correlation within a genotype between Z-type female behavior and male courtship that facilitates choice by another Z-type female genotype (HOLLOCHER *et al.* 1997 ), suggesting that these strains could be classified as having Z-type male behavior. The sets of strains include some used to originally describe the Z/M system collected in the early 1990s (WU *et al.* 1995; HOLLOCHER *et al.* 1997) and another set collected in the 2010s. Strains Z53 and Z30 are considered as "strong Z" (WU *et al.* 1995, provided by Trudy Mackay) and strain Z29 as "moderate" Z (WU *et al.* 1995; HOLLOCHER *et al.* 1997, BDSC #60741). Strains ZH33, ZH42 (provided by Andy Clark, GRENIER *et al.* 2015) and ZS11 (provided by Chip Aquadro) have previously been phenotyped for female Z behavior (HOLLOCHER *et al.* 1997). Strains ZK82 and ZK58 were collected in Lake Kariba, Zimbabwe but not previously phenotyped (provided by Chip Aquadro). Lower Zambize 2.1 (LZ21), Chipata 11 (CH11), Chipata 12 (CH12),

Livingstone 4.7 (LS47), and Lusaka Camp 4 (LC4) were collected in Zambia in 2015 by Daniel Matute and are all Z-type (Matute personal communication).

***Quantification of CHCs***

We ran samples on a Shimadzu GC 2014 with flame ionization detector with an Agilent DB-5 column (20m long, 0.18mm inner diameter, 0.18μm film thickness). We injected 2μl of sample into a preheated injection port at 290°C using the AOC-20i autoinjector. After injection the sample was equilibrated at 50°C for 1 min. The oven temperature was increased to 150°C at a rate of 20°C/minute, and then ramped to 300°C at 5°C/min followed by a final hold at 300°C for 10 minutes. All samples were run in splitless mode, with a sampling time of 1 min and split ratio of 20, helium as the carrier and makeup gas, a constant linear velocity of 35cm/sec, and purge flow of 3ml/min split. The detector temperature was 340°C and sampling rate was 40cm/sec. The compounds were identified based on Kovats retention index, which compares retention time of known and unknown compounds, and compared to previous reports for CHCs in *D. melanogaster* that captured presumptive Z-type compounds (DEMBECK *et al.* 2015). The area of each peak was divided by the area of the hexacosane (C27) internal standard, so quantities could be compared between samples.

***Statistical analysis***

*Comparison of courtship between recording periods*

To ensure that the same behaviors we observed in our initial cataloging (Fall 2018) were consistent with behaviors when determining selection (Spring 2019) we compared common genotype combinations between these two collection periods and analyzed Z53 female x Z53 female and DGRP882 female x DGRP882 male interactions. We first looked at clustering of these observations using principal component analysis. Besides a single outlier for DGRP882 in

2018, the genotypes were clearly distinct with no difference between years (Supplemental Fig. 1). Specifically DGRP882 sang more in both years compared to Z53, and the rate of singing was the same for both recording periods (Supplemental Fig. 2). Similarly Z53 always scissored more compared to DGRP882, and this behavior was consistent across years (Supplemental Fig 2).

*Quantification of courtship effort (courtship index) in trials that failed to copulate*

We quantified behavior for interactions that ended in copulation because these were the only useful trials to make correlations between phenotypes and copulation latency (fitness). However, we wanted to ensure that we were justified in not using trials where no copulation occurred as representative of the behavioral sequence. Additionally because no copulations occurred between Z53 females and DGRP882 males we needed to ensure that these males were actively courting to be able to include them in our analyses. We found that there was no difference in the time spent courting for DGRP882 female x DGRP882 male and Z53 female x DGRP882 male trials, indicating a high courtship index (Wilcox Rank sum test, $W = 16$, p-value $= 0.9158$). The average courtship index for DGRP882 female x DGRP882 male was 0.98 and the average courtship for Z53 female x DGRP882 male was 0.96. This indicated males were actively courting and females were rejecting them.

To determine if males that failed to copulate were courting in other genotype combinations we compared successful and non-successful matings between Z29 males and either DGRP882 females or Z53 females. We chose the Z29 male genotype because there was an equal proportion of males that did and did not successfully copulate with both female genotypes. While the small sample size precluded formal statistical analysis, the patterns between trials that ended in copulation compared to trials that did not were very obvious (Supplemental Fig 3). The average courtship index for DGRP882 female x Z29 male trials that copulated was 0.94, compared to a courtship index of 0.54 for trials that did not copulate. Similarly, the average courtship index for Z53 female x Z29 male trials that copulated was 0.92 compared to a courtship index of 0.44 for trials that did not copulate. From this comparison we were confident the majority of trials that failed to copulate were the result of males not courting and could be excluded from the analyses. We did not determine whether courtship was initially strong, and later tapered off due to female cues/female rejection, or if these males were deficient in courtship, but it is typically thought that *D. melanogaster* males vigorously court, even outside of their own species (BARKER 1967; CARRACEDO *et al.* 2000).

*Mating rejection in long-term experiments*

After analyzing videos we observed several genotype combinations that failed to copulate. We expected Z53 to reject known M-type males, and consistent with this prediction we did not observe copulations between Z53 females and DGRP882 or Canton S males. There were several genotype combinations that we did not a priori expect to fail including Z53 females x LZ21 males and DGRP882 females with Z30 or LZ21 males. To determine if this was a function of our short recording time we set up single pair matings between these genotypes and measured the time until progeny were first seen in vials over a 16 day period. We included DGRP882 x DGRP882 and Z53 x Z53 as controls. We also included Z53 female x ZH33 male and DGRP882 female x ZH33 male as representative genotype combinations that mated at intermediate levels with both female genotypes. We initially set up eight single pair crosses per genotype combination and examined these crosses daily for the presence of eggs and/or larvae for 16 days. During the course of the experiment we had many females and males die before any eggs were laid and excluded these from further analyses. We compared the number of days until we observed the first egg or first larvae by fitting survival models for each female genotype separately. We fit Cox proportional hazards models using the survival package in R (THERNEAU 2020) because these models can take into account observations that were stopped (censored) at the end of the experiment. These models conveniently summarize the number of crosses where we observe eggs/larvae compared to the number of initial crosses over time. At the end of 16 days if a cross had not produced eggs or larvae and both individuals were still alive, we considered this observation to be censored.

*VSURF variable reduction and ART ANOVA for plasticity models*

Because the transition matrix was large and many transitions were correlated, we used the VSURF package in R (GENUER *et al.* 2019) for variable reduction. we set up a model where transition frequency could be explained by a categorical variable that captured the interaction of male x female genotype, which is essentially a classification problem. VSURF uses thresholds to determine variables it will evaluate, interpretation to identify variables that strongly contribute to the dependent variable, and then a prediction step to reduce the number of correlated variables. From the remaining variables we carried out ART ANOVA testing for male, female and male x female genotype effects using the ARTools package for R (KAY AND WOBBROCK 2019). These models are non-parametric which was well suited for our analysis since the behavioral variables were bound by 0-1, did not produce normally distributed residuals in standard ANOVAs, and had large differences in variance between groups (heteroskedasticity). All three characteristics would have made standard ANOVA inappropriate (FOX 2016). We favored this non-parametric approach instead of transforming the data since not all transformations maintain additivity and constant variance (MCULLAGH AND NELDER 1989; FENG *et al.* 2014). Instead of transformation we chose to use quantile regression which is robust to these issues (KOENKER 2005).

**Supplemental Table 1.** Male courtship behaviors are not highly correlated and are not consistently correlated when males court either female genotype. Pairwise correlations were calculated for trials where males courted DGRP882 females (above diagonal) and trials where males courted Z53 females (below diagonal).

| | Engaging | Attempting | Singing | Singing-2 | Circling | Scissoring |
|---|---|---|---|---|---|---|
| Engaging | | -0.13 | -0.69 | 0.00 | -0.09 | -0.09 |
| Attempting | -0.40 | | -0.16 | -0.26 | -0.13 | -0.09 |
| Singing | -0.38 | -0.30 | | 0.24 | -0.02 | -0.30 |
| Singing-2 | -0.18 | -0.23 | 0.41 | | 0.32 | -0.30 |
| Circling | -0.23 | 0.08 | -0.24 | 0.04 | | -0.12 |
| Scissoring | -0.04 | -0.48 | -0.13 | -0.05 | -0.08 | |

**Supplemental Table 2.** Male courtship behavioral traits are female-genotype dependent. The male genotype effect captures differences between male genotypes in a courtship trait. The female genotype effect indicates plasticity. An interaction effect indicates that changes in male genotype behavior is not consistently parallel across genotypes. Significance was determined using ART ANOVA and significant effects (*P*<0.05) are reported in bold.

| | Male genotype effect | | Female genotype effect | | Interaction | |
|---|---|---|---|---|---|---|
| Trait | F value | P-value | F value | P-value | F value | P-value |
| Engaging | **5.05** | **<0.0001** | 0.031 | 0.8598 | **2.264** | **0.0211** |
| Attempting | **3.88** | **0.0002** | **28.844** | **<0.0001** | 1.759 | 0.0808 |
| Singing | **7.93** | **<0.0001** | **15.826** | **0.0001** | 1.016 | 0.4363 |
| Circling | **3.92** | **0.0002** | **29.846** | **<0.0001** | **4.073** | **0.0001** |
| Scissoring | **8.49** | **<0.0001** | 3.743 | 0.0563 | 1.596 | 0.1214 |

**Supplemental Table 3.** Male courtship plasticity changes the transition matrix for some, but not all male genotypes. Homogeneity tests compared the complete transition matrix describing the behavioral sequence when males courted Z-type and M-type females.

| Male genotype | \|2 statistic | Degree of Freedom | *P*-value |
|---|---|---|---|
| Z53 | 84.084 | 35 | <0.001 |
| Z29 | 80.839 | 35 | <0.001 |
| ZH33 | 163.217 | 35 | <0.001 |
| ZH42 | 11.512 | 35 | 0.999 |
| LC | 15.610 | 35 | 0.998 |
| LS4-7 | 185.410 | 35 | <0.001 |
| ZK58 | 144.166 | 35 | <0.001 |
| ZK82 | 110.480 | 35 | <0.001 |
| CH11 | 25.423 | 35 | 0.882 |
| CH12 | 46.36 | 35 | 0.094 |
| ZS11 | 108.463 | 35 | <0.001 |

**Supplemental Table 4.** Male courtship transition plasticity is female-genotype dependent. The male genotype effect captures differences between male genotypes in a courtship trait. The female genotype effect indicates plasticity. An interaction effect indicates that changes in male genotype behavior is not consistently parallel across genotypes. Significance was determined using ART ANOVA and significant effects ($P<0.05$) are reported in bold.

| | Male genotype effect | | Female genotype effect | | Interaction | |
|---|---|---|---|---|---|---|
| Transition | F value | P-value | F value | P-value | F value | P-value |
| Sing-Scissor | **6.55** | **<0.0001** | 3.08 | 0.0827 | 0.61 | 0.7995 |
| Sing-Attempt | **3.48** | **<0.0001** | **16.78** | **<0.0001** | 1.41 | 0.1871 |
| Eng-Sing | **2.38** | **0.0151498** | **9.62** | **0.0026** | 0.51 | 0.8736 |

**Supplemental Table 5** Male cuticular hydrocarbon (CHC) plasticity is female-genotype dependent. The male genotype effect captures differences between male genotypes in a courtship trait. The female genotype effect indicates plasticity. An interaction effect indicates that changes in male genotype behavior are not consistently parallel across genotypes. Significance was determined using ART ANOVA and significant effects (*P*<0.05) are reported in bold.

| | Male genotype | | Female genotype | | Interaction | |
|---|---|---|---|---|---|---|
| CHC | F value | P-value | F value | P-value | F value | P-value |
| 5-C25 | **47.08** | **<0.0001** | **4.96** | **0.0192** | 1.55 | 0.2376 |
| 9-C25 | **48.18** | **<0.0001** | 2.76 | 0.0895 | 1.92 | 0.1740 |
| 7-C23 | **8.96** | **0.0077** | 0.97 | 0.3965 | **4.30** | **0.0296** |
| n-C22 | 1.04 | 0.3197 | 2.87 | 0.0822 | 0.005 | 0.9950 |
| 2-Me-C30 | **32.45** | **<0.0001** | 0.005 | 0.9943 | **6.76** | **0.0064** |

**Supplemental Table 6.** Directional selection on male courtship behavior is female-genotype dependent. Selection coefficients for male courtship traits and mating success, measured by copulation latency, demonstrate which traits are under selection. The intercept is determined for the DGRP-882 genotype. Slope M represents the selection coefficient for the M genotype DGRP-882. The Z effect is the change in intercept for the Z-type Z53. The Slope change Z is the change in the selection gradient in the Z type Z53 strain. The confidence intervals were fit using quantile regression. Confidence intervals that do not overlap zero are reported in bold.

| | Intercept | | Slope M | | Z effect | | Slope change Z | |
|---|---|---|---|---|---|---|---|---|
| Trait | Lower | Upper | Lower | Upper | Lower | Upper | Lower | Upper |
| Engaging | **5.74** | **19.73** | -38.73 | 56.68 | -8.913 | 9.92 | -79.86 | 19.905 |
| Attempting | -7.62 | 14.08 | **7.90** | **109.91** | **2.00** | **26.17** | **-138.14** | **-29.55** |
| Singing | **14.65** | **31.65** | -40.70 | 19.89 | **-36.40** | **-11.11** | **26.27** | **59.41** |
| Circling | **7.71** | **19.34** | -198.91 | 150.41 | -17.77 | 5.83 | -197.32 | 244.54 |
| Scissoring | **4.39** | **9.53** | **35.24** | **60.46** | -4.69 | 3.28 | -25.62 | 29.97 |
| Separate | **7.31** | **16.55** | -76.75 | 23.11 | -10.21 | 3.02 | -84.41 | 160.77 |

**Supplemental Table 7.** The probability that mating occurs based on time spent exhibiting specific behaviors is female-genotype dependent. The intercept is determined for the DGRP882 genotype. OR M represents the estimate for the Odds Ratio for the M genotype DGRP882. The Z effect is the change in intercept for the Z-type Z53. The OR change Z is the change in the odds ratio in the Z type Z53 strain. The confidence intervals were fit using binomial regression. Confidence intervals that do not overlap zero are reported in bold.

| | Intercept | | OR M | | Z effect | | OR change Z | |
|---|---|---|---|---|---|---|---|---|
| Trait | Lower | Upper | Lower | Upper | Lower | Upper | Lower | Upper |
| Singing | **-6.17** | **-1.58** | **4.13** | **14.41** | **3.27** | **8.91** | **-20.80** | **-7.37** |
| Scissoring | **0.06** | **1.74** | -11.74 | 0.34 | -2.18 | 0.22 | **1.45** | **20.66** |
| Circling | -0.25 | 1.22 | -35.44 | 14.29 | -0.90 | 1.19 | -19.33 | 34.80 |

**Supplemental Table 8.** Directional selection on male cuticular hydrocarbons (CHCs) is female-genotype dependent. Selection coefficients for male CHCs and mating success, measured by copulation latency, demonstrate which CHCs are under selection. Selection coefficients representing relationships between male cuticular hydrocarbons (CHCs) and mating success, measured by copulation latency. The intercept is determined for the DGRP-882 genotype. Slope M represents the selection coefficient for the M genotype DGRP-882. The Z effect is the change in intercept for the Z-type Z53. The Slope change Z is the change in the selection gradient in the Z type Z53 strain. The confidence intervals were fit using quantile regression. Confidence intervals that do not overlap zero are reported in bold.

| | Intercept | | Slope M | | Z effect | | Slope change Z | |
|---|---|---|---|---|---|---|---|---|
| CHC | Lower | Upper | Lower | Upper | Lower | Upper | Lower | Upper |
| n-C24 | **1.22** | **6.40** | **555.26** | **820.43** | **3.18** | **15.53** | **-1427.02** | **-473.52** |
| 6,10-C25 | **3.90** | **18.08** | -136.20 | 351.52 | -9.57 | 8.27 | -242.37 | 92.40 |
| 9-C25 | **1.03** | **12.02** | **14.39** | **421.64** | -2.66 | 19.48 | **-470.39** | **-95.24** |
| n-C29 | **5.92** | **7.28** | -84.71 | 85.92 | -0.64 | 1.09 | -203.46 | 41.94 |
| 2-Me-C26 | -11.80 | 4.51 | **63.97** | **181.45** | **1.24** | **20.30** | **-199.78** | **-10.53** |
| 7-C23 | **1.91** | **27.62** | -32.73 | 12.96 | -24.97 | 8.13 | -35.10 | 42.51 |
| cVA | **11.63** | **18.55** | **-74.19** | **-1.09** | **-12.35** | **-4.24** | **48.15** | **155.92** |

**Supplemental Table 9.** The probability that mating occurs based on cuticular hydrocarbon (CHC) amount is female-genotype dependent. The intercept is determined for the DGRP-882 genotype. Slope M represents the estimate for the Odds Ratio for the M genotype DGRP-882. The Z effect is the change in intercept for the Z-type Z53. The slope change Z is the change in the odds ratio in the Z type Z53 strain. The confidence intervals were fit using binomial regression. Confidence intervals that do not overlap zero are reported in bold.

| | Intercept | | Slope M | | Z effect | | Slope change Z | |
|---|---|---|---|---|---|---|---|---|
| CHC | Lower | Upper | Lower | Upper | Lower | Upper | Lower | Upper |
| 7-C23 | -2.31 | 0.83 | -1.54 | 4.81 | 0.65 | 5.37 | **-10.37** | **-0.95** |
| 5-C23 | -0.36 | 1.94 | -24.52 | 4.51 | -2.94 | 0.39 | -0.42 | 41.70 |
| n-C23 | -2.15 | 0.73 | -4.27 | 14.59 | **0.57** | **4.77** | **-29.84** | **-2.65** |
| cVA | **-1.90** | **-0.04** | **2.23** | **22.22** | **0.82** | **3.49** | **-36.68** | **-8.41** |

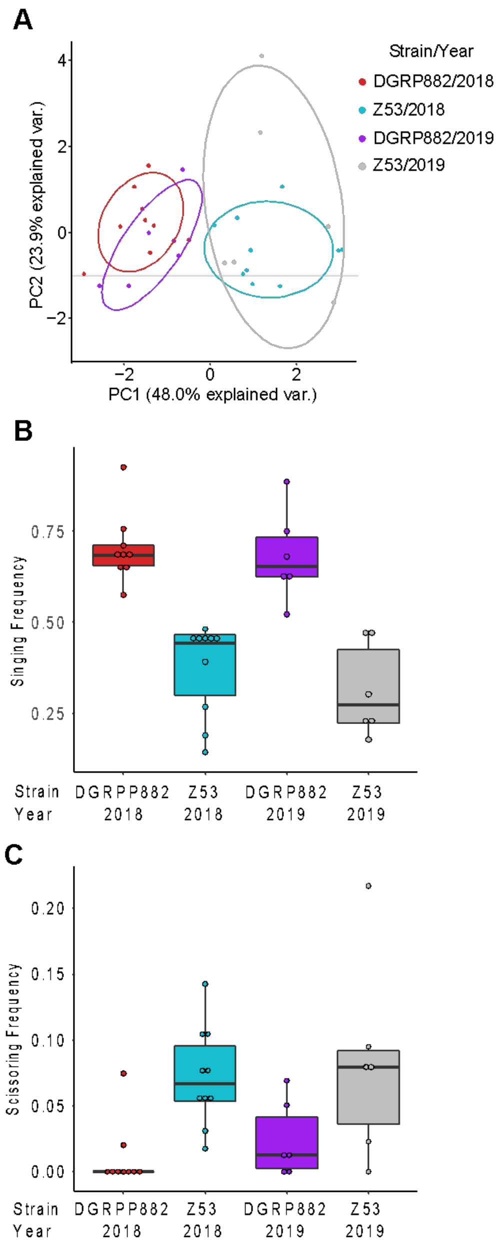


**Supplemental Figure 1.** Male courtship behaviors are consistent between years with discernable differences between DGRP-882 and Z53 males in both years. A) Principal component analysis (PCA) of behavioral frequencies demonstrates separation of the male genotypes. B) In both years DGRP-882 males have significantly higher Singing frequency compared to Z53 males. C) In both years Z53 males have significantly higher scissoring frequency compared to DGRP-882 males.

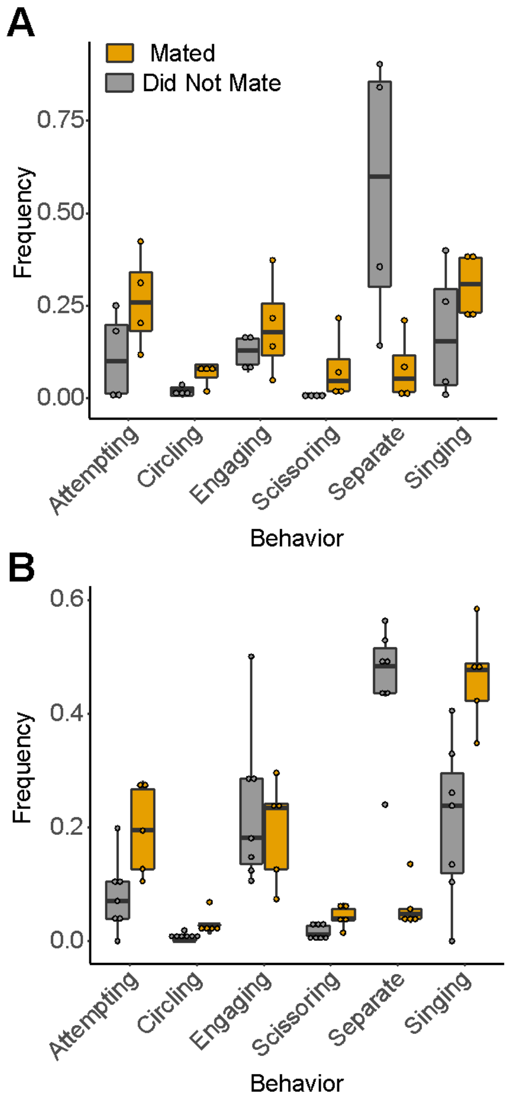


**Supplemental Figure 2.** Males that did not copulate (mate) failed to court vigorously. Z29 males that interacted with either A) Z53 females or B) DGRP-882 females copulated when actively courting. Males that did not copulate courted less actively, indicated by the higher amount of time spent "separate". In each panel, the data were separated into two groups depending on whether mating occurred.

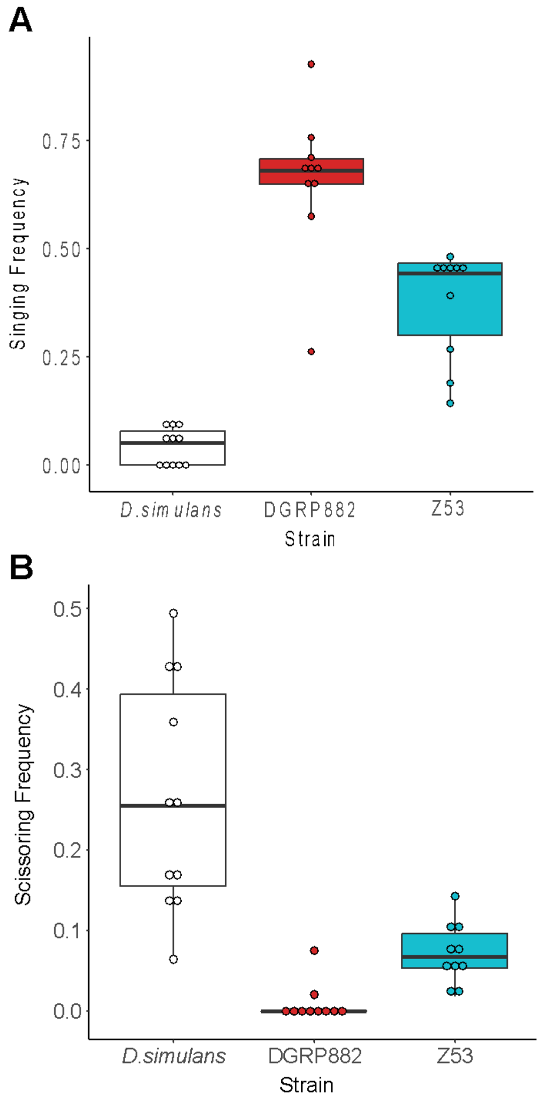


**Supplemental Figure 3**. Z-type males exhibit differences in key behaviors compared to both M-type *D. melanogaster* and *D. simulans,* including both A) singing and B) scissoring frequencies. A) There were significant differences in singing frequencies with DGRP-882 males exhibiting the highest frequencies and *D. simulans* the lowest. B) There were significant differences in scissoring frequencies with *D. simulans* exhibiting the highest and DGRP-882 the lowest. Most of the DGRP-882 males exhibited close to zero scissoring behavior.

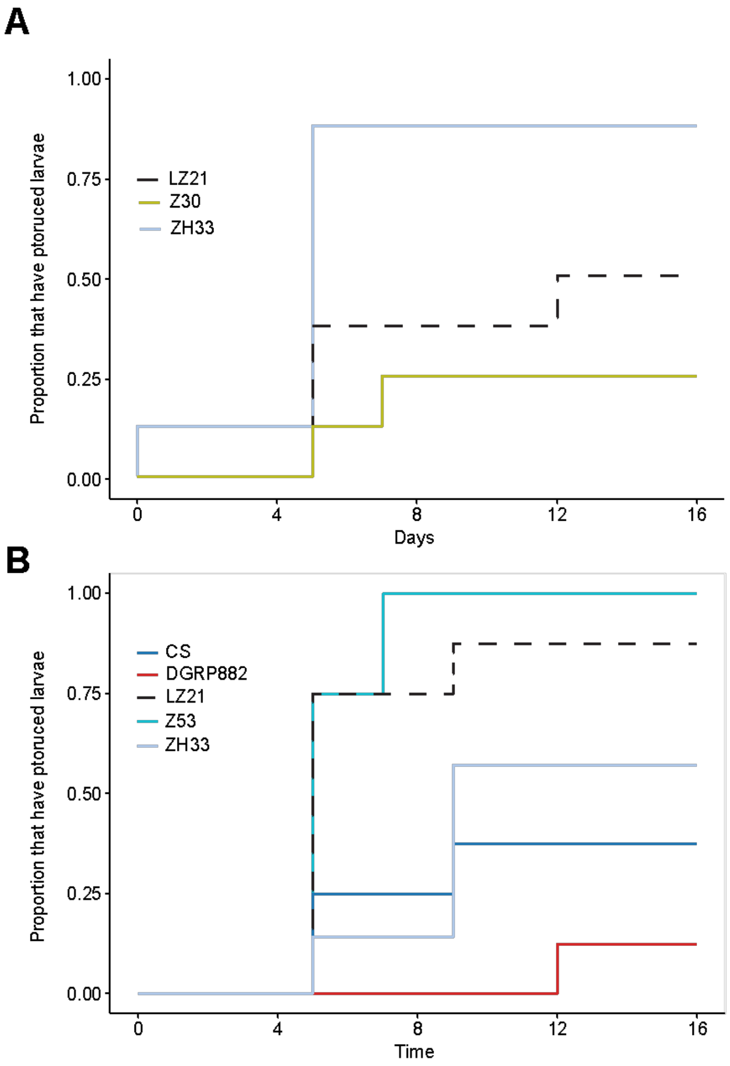


**Supplemental Figure 4**. Reproductive isolation between M-type and Z-type strains. A) Male genotypes were paired with DGRP882 females and showed significant variation in the number that mated, with mating status inferred by the presence of larvae. The X axis shows the time since flies were paired. Pairs that mated within the first two days had larvae present around day 4-5, such as ZH33. The remaining genotypes took longer to mate with some males not mating even after 16 days. B) Male genotypes were paired with Z53 females and showed significant variation in the number that mated and days until mating as inferred by the presence of larvae.

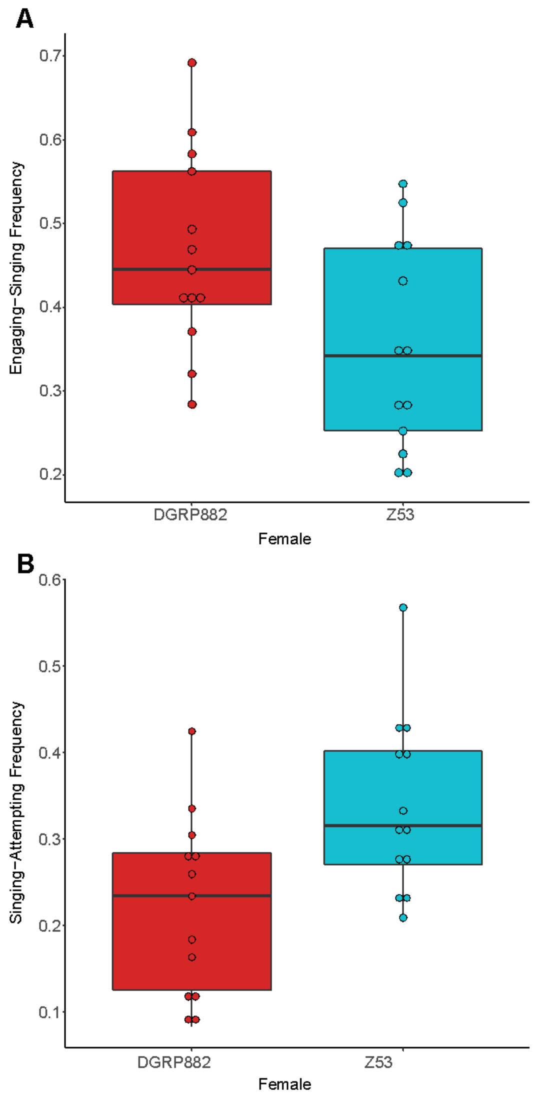


**Supplemental Figure 5.** Plasticity in male behavioral frequencies results in differences in the transitions between behavioral states. A) The engaging-singing transition frequency was significantly higher when a male was presented to a DGRP-882 female than to a Z53 female, consistent with more singing when males interacted with DGRP-882 females. B) The singing-attempted copulation transition frequency was significantly lower when a male was presented to a DGRP-882 female than to a Z53 female, consistent with more attempting and less singing when males interacted with Z53 females. Each point in the figures represents the average frequency of the corresponding transition of all males in one genotype, when presented to the given female.

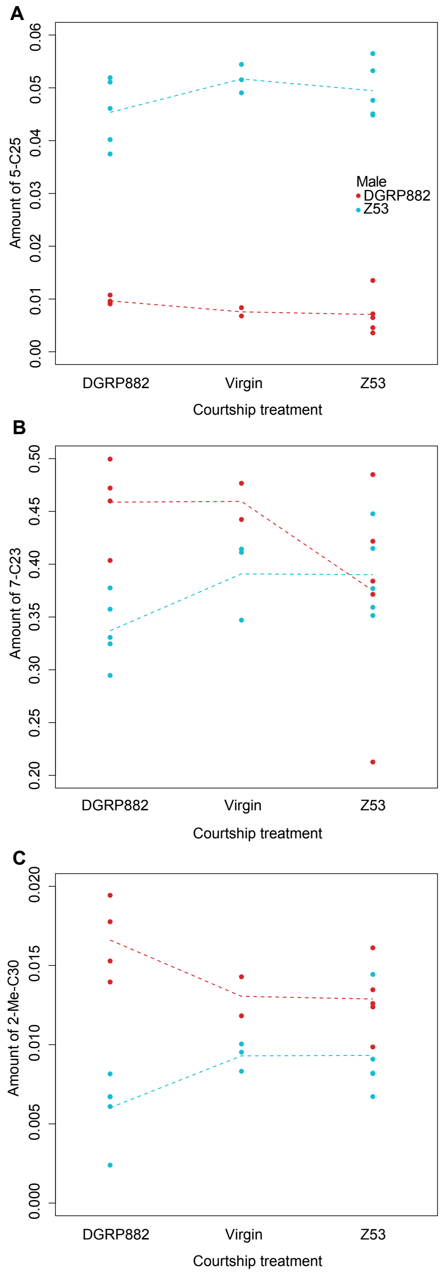


**Supplemental Figure 6.** Male cuticular hydrocarbons are plastic and change differently for the male genotypes based on mating exposure. A) The amount of 5-C25 produced by Z53 males decreased when males were presented to females of either genotype. B) The amount of 7-C23 produced by each male genotype remained the same when females of the same genotype were presented to it, but decreased when the females had a different genotype. C) The amount of 2-Me-C30 changes in opposite directions when males were presented with DGRP882 females. Each dot represents an individual male. Lines are the average amount for each treatment.

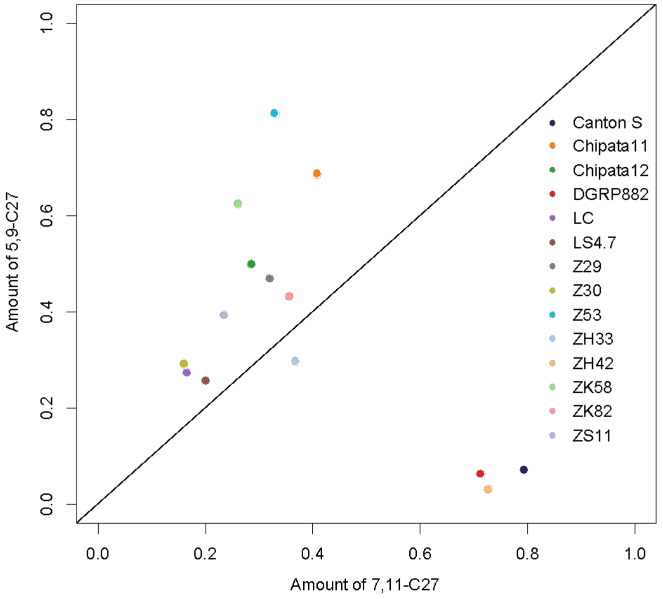


**Supplemental Figure 7.** The relationship between the two major female cuticular hydrocarbons (CHCs) that have historically defined M and Z-types. There is no overall tradeoff between the amount of 7,11-heptacosadiene (7,11-C27) and 5,9-heptacosadiene (5,9-C27), instead there is an approximately bimodal distribution. The known M-type females DGRP-882 and Canton S have high levels of 7,11-C27 and almost no 5,9-C27, whereas females collected from regions in Southern Africa have different ratios of these two compounds.

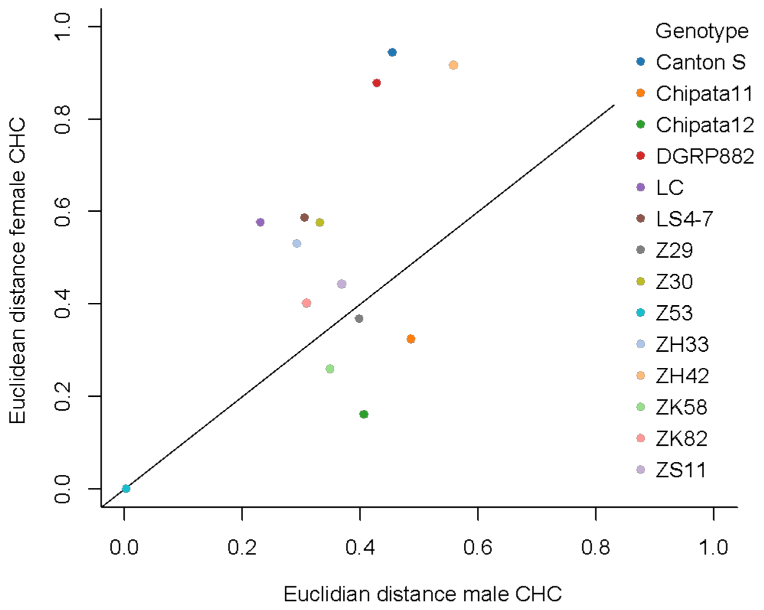


**Supplemental Figure 8.** Male and female CHC profiles are correlated. The Euclidean distance was calculated with respect to the Z53 genotype, so Z53 is centered at the origin. The M-type strains, DGRP882 and Cantons S, and ZH42 are divergent from Z53 for both male and female CHCs. The remaining Z-type genotypes have large variation in CHC profiles with some genotypes divergent from both Z53 and the M-type strains.

**Supplemental Video 1.** An example of the circling behavior observed in Z-type strains.

**Supplemental Video 2.** An example of the scissoring behavior observed in Z-type strains.

**Supplemental Video 3.** An example of the scissoring behavior observed in *D. simulans*.
